# A databank for intracellular electrophysiological mapping of the adult somatosensory cortex

**DOI:** 10.1101/472043

**Authors:** Angelica da Silva Lantyer, Niccolò Calcini, Ate Bijlsma, Koen Kole, Melanie Emmelkamp, Manon Peeters, Wim J. J. Scheenen, Fleur Zeldenrust, Tansu Celikel

## Abstract

**Background:** Neurons in the supragranular layers of the somatosensory cortex integrate sensory (bottom-up) and cognitive/perceptual (top-down) information as they orchestrate communication across cortical columns. It has been inferred, based on intracellular recordings from juvenile animals, that supragranular neurons are electrically mature by the fourth postnatal week. However, the dynamics of the neuronal integration in the adulthood is largely unknown. Electrophysiological characterization of the active properties of these neurons throughout adulthood will help to address the biophysical and computational principles of the neuronal integration.

**Findings:** Here we provide a database of whole-cell intracellular recordings from 315 neurons located in the supragranular layers (L2/3) of the primary somatosensory cortex in adult mice (9-45 weeks old) from both sexes (females, N=195; males, N=120). Data include 361 somatic current-clamp (CC) and 476 voltage-clamp (VC) experiments, recorded using a step-and-hold protocol (CC, N=257; VC, N=46), frozen noise injections (CC, N=104) and triangular voltage sweeps (VC, 10 (N=132), 50 (N=146) and 100 ms (N=152)), from regular spiking (N=169) and fast-spiking neurons (N=66).

**Conclusions:** The data can be used to systematically study the properties of somatic integration, and the principles of action potential generation across sexes and across electrically characterized neuronal classes in adulthood. Understanding the principles of the somatic transformation of postsynaptic potentials into action potentials will shed light onto the computational principles of intracellular information transfer in single neurons and information processing in neuronal networks, helping to recreate neuronal functions in artificial systems.

## Data description

The primary somatosensory cortex (S1) encodes time-varying but spatially well defined haptic information [1] from the mechanoreceptors in the skin, thereby creating a topographical neuronal representation of the tactile world [2, 3]. Rodents, for example, locate tactile targets in their immediate environment by integrating information across these (whisker) representations in the barrel cortex [4], where neurons in each cortical column preferably respond to a single whisker on the contralateral snout [5]. The supragranular layers (cortical layers 2/3, L2/3) of the barrel cortex is the first cortical network that integrates the sensory information across neighboring cortical columns, whiskers and whisk cycles [6–10]. This representation of the whisker contacts undergoes experience-dependent changes [11–14] and is altered in animal models of neurodevelopmental disorders [15–17]. Adaptive changes in the synaptic and modulatory drive could powerfully regulate the transformation of postsynaptic responses into action potentials, ultimately controlling how sensory information is transferred between cortical columns and cortical regions [18].

Understanding the principles of neuronal information transfer in the supragranular layers will require a systematic analysis of the integrative properties of these cortical neurons. Thus far, however, slice experiments primarily focused on juvenile animals as it is widely considered that the neurons mature anatomically and electrophysiologically by the fourth postnatal week [16, 19–23]. Here we provide a database of 837 experiments collected from 315 adult supragranular neurons that will help to address the principles of information processing by cortical neurons throughout the adulthood of mice. The database consists of whole-cell intracellular recordings in voltage-clamp (VC) and current-clamp (CC) configurations: while current-clamp somatic measurements bring insight into the properties related to action potential initiation, timing, rate, and pattern, voltage-clamp recordings provide information on the voltage-gated ion-channel dynamics. The database is best utilized to address the principles of information transfer in individual neurons (see e.g. [18, 24]) and for the electrical characterization of adult cortical sensory neurons. It will serve synaptic, systems, computational and theoretical neuroscientist in search of the principles of information processing, transfer, and recovery in neuronal networks. The database is expected to create synergy with 1) the recently completed transcriptome [25, 26] and proteome [27, 28] of the supragranular layers of the barrel cortex, 2) computational models of the molecular changes that contribute to the maturation of synaptic communication in the same cortical region (e.g [29]), 3) computational models of synaptic integration and action potential generation in the supragranular layers of the barrel cortex [18], and 4) the high-resolution mapping of sensory representations using intrinsic signals in single trial resolution (e.g. [30]) resulting in a multi-scale analysis of the cortical organization, from molecules of chemical communication to network representations.

### Methods

Experiments that involve animals were conducted in accordance with the European Directive 2010/63/EU, national regulations in the Netherlands, and international guidelines on animal care and use of animals. Pvalbtm1(cre)Arbr (RRID:MGI:5315557) or Ssttm2.1(cre)Zjh/J mice (RRID:IMSR_JAX:013044) from either sex (N=75 females, N=45 males, aged 9-45 weeks) were used from the local breeding colonies.

The mice were anesthetized with Isoflurane (1.5 ml/mouse) before the tissue was extracted and coronal slices of the primary somatosensory cortex, barrel subfield region is prepared (Figure 1). The procedures were as described before [11, 12, 14, 16, 31] with the exception that animals were intracardially perfused with ice-cold dissection solution containing (in mM) 108 choline chloride, 3 KCl, 26 NaHCO_3_, 1.25 NaH_2_PO_4_.H_2_O, 25 Glucose.H_2_O, 1 CaCl_2_.2H_2_O, 6 MgSO_4_.7H_2_O, 3 sodium pyruvate, after animals were deeply anesthetized, as assessed by pinch withdrawal reflex, heart and breathing rate. The brain was removed after decapitation and sliced coronally (300 micrometers in thickness) in the same ice-cold perfusion medium. The slices were then transferred to a chamber containing aCSF (in mM): 120 NaCl, 3.5 KCl, 10 Glucose.H_2_O, 2.5 CaCl_2_.2H_2_O, 1.3 MgSO_4_.7H_2_O, 25 NaHCO_3_, 1.25 NaH_2_PO_4_.H_2_O, aerated with 95% O_2_/ 5% CO_2_ at 37^□^C. After 30 minutes, the slices were transferred to room temperature before whole-cell electrophysiological recordings started.

**Figure 1.**
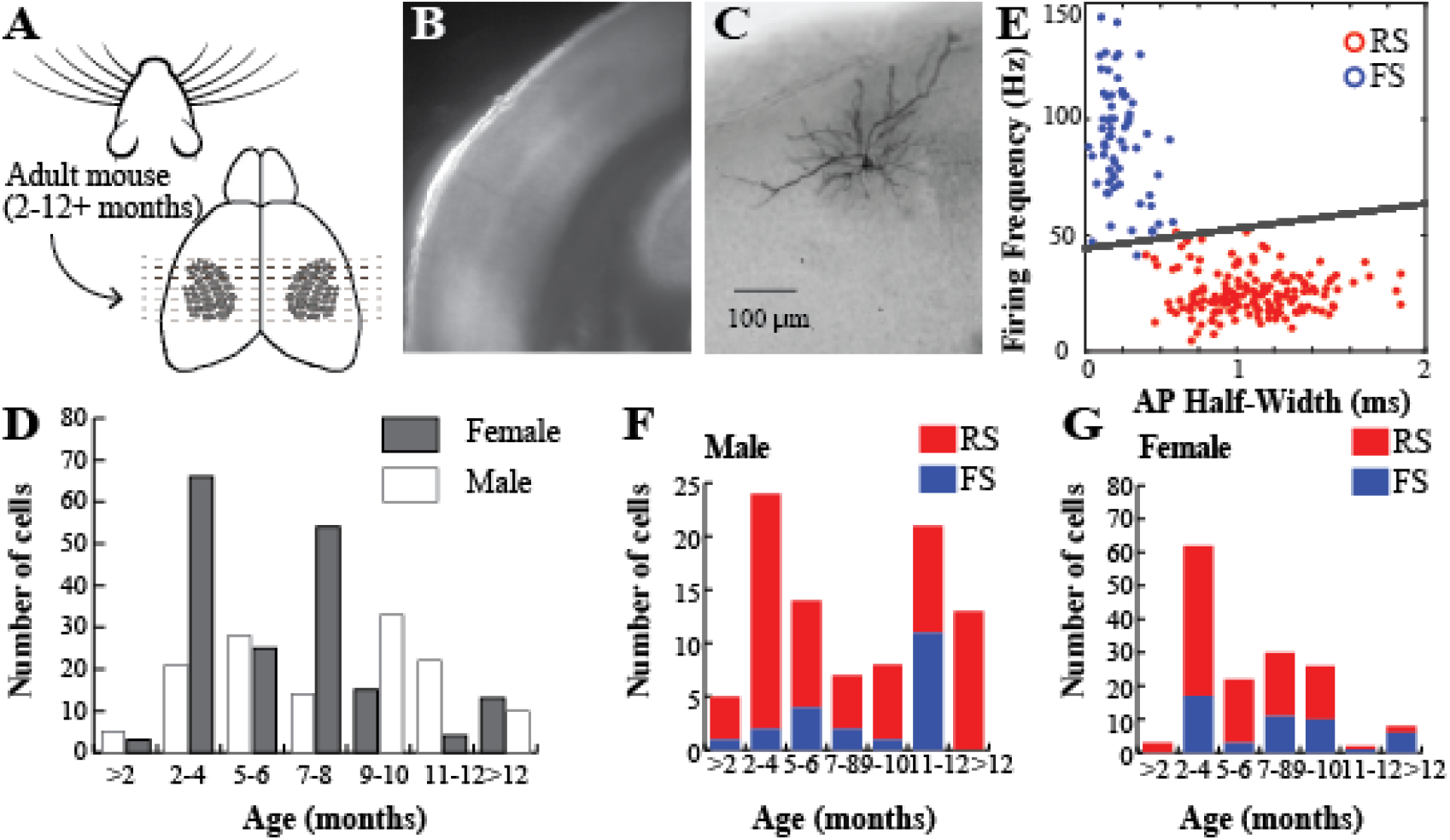
Acute slice preparation. **(A)** Coronal slices (300 micrometers in thickness) were prepared for ex vivo recording from the L2/3 neurons in the mouse primary somatosensory cortex, barrel cortex subregion. **(B)** A low magnification view of the slice in 4x. **(C)** A representative neuron, intracellularly filled with biocytin and visualized with DAB staining (VECTASTAIN Elite ABC Kit, RRID: AB_2336827) according to the manufacturer’s guidelines. **(D)** Distribution of the 326 neurons in this database across males (39.9%) and females (60.1%) as well as the ages of the animals. **(E)** Classification of the neurons presumed fast-spiking (FS) and regular spiking (RS) populations based on firing frequency and action potential half-width (see Methods for details). **(F-G)**. The distribution of cells across cell type and ages.

### Whole-cell recordings

Slices were continuously oxygenated and perfused with aCSF during recordings. The barrel cortex was localized and cells of interest in the supragranular layers were patched under 40x magnification in room temperature using HEKA EPC 9 and EPC10 amplifiers in combination with the Patch Master v2×90.2 data acquisition software. The data band-pass filtered (0.1- 3000Hz). The AC mains (hum) noise (max peak-to-peak amplitude 0.2 mV) that exists in a subset (~4%) of the recordings was not filtered. Patch clamp electrodes were pulled from glass capillaries (1.00 mm (external diameter), 0.50 mm (internal diameter), 75 mm (length), GC100FS-7.5, Harvard Apparatus) with a P-2000 puller (Sutter Instrument, USA) and used if their initial resistance were between 5 and 10 MOhm. They were filled with intracellular solution containing (in mM) 130 K-Gluconate, 5 KCl, 1.5 MgCl_2_.6H_2_O, 0.4 Na3GTP, 4 Na2ATP, 10 HEPES, 10 Na-phosphocreatine, 0.6 EGTA, and the pH was set at 7.22 with KOH. Current-clamp and voltage-clamp recordings were performed as described before [32], [33] and included four stimulus protocols (Figure 2).

**Figure 2.**
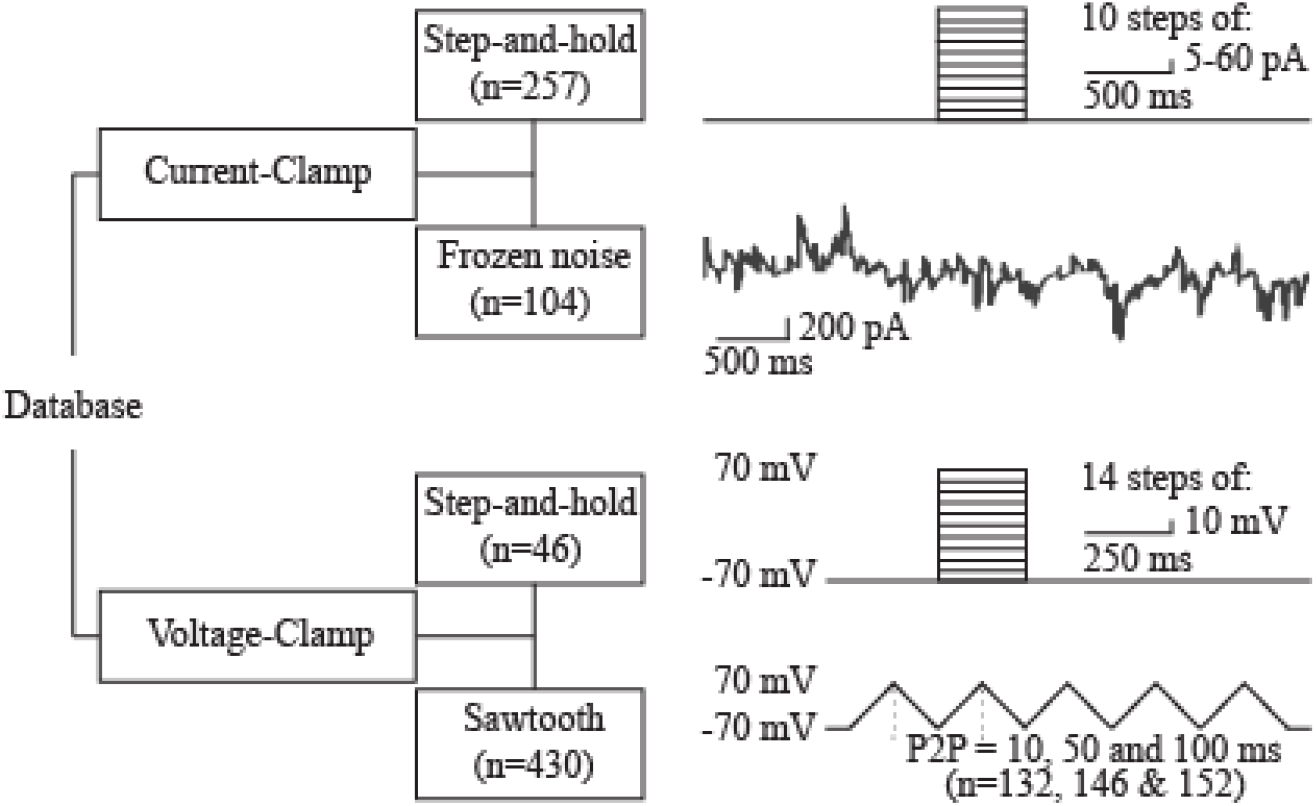
Experimental protocols and the hierarchical organization of the database. The data is available online at http://gigadb.org/dataset/100535. The database contains two subfolders, current-clamp and voltage-clamp, each of which has additional subfolders based on the stimulus protocols utilized in this study. Each dataset is provided in a .mat format and includes both voltage and current channels unless otherwise described. The stimulus delivered to the cells as well as the cell’s response can be quantified from these variables.

### Current-clamp protocol

After establishing the current-clamp configuration, the resting membrane potential was set to −70 mV by direct somatic current injections, as required. The step-and-hold stimulation protocol included 10 steps of 500 ms long depolarization pulses (step size: 5, 10, 20, 40 or 60 pA) with an inter-sweep-interval of 6.5 s. The stimulus train was repeated 1-3 times with a 20 s interval. The drift, if any, in resting membrane potential during the recording was not corrected for. However, any neuron whose resting membrane potential varied more than 7 mV was not included in the database. The frozen-noise (FN) stimulation protocol involved somatic injection of the current that is the output of an artificial neural network of 1000 neurons, each firing Poisson spike trains in response to a ‘hidden state’ (see [34] for details, and [35] on how to generate the frozen noise input and analyze the data).

### Voltage-clamp protocol

The voltage-clamp stimulation protocols included step-and-hold and sawtooth (triangular) pulse injections (Figure 2). In both protocols, the membrane potential was clamped at −70 mV prior to somatic depolarization. In the step-and-hold protocols, 14 incremental steps of depolarizing pulses (10 mV/each) were delivered for a period of 250 ms with an interval of 20 s. Sawtooth pulses (range: −70 to 70 mV) were delivered at three frequencies (5, 10, 50 Hz) and consisted of five triangular pulses with peak-to-peak (P2P) distances of 200, 100, 20 ms, respectively. Each trial was repeated twice with 20 s interval.

### Data organization

Files in “.mat” (MATLAB) format containing the original traces from each experiment are organized in folders separated by the structure described in Figure 2. Metadata including the date and number of the experiments, the experimenter’s initials, the animal’s sex and age,the experimental protocol, the cell type, and the animal number are included in a tabulated format (.xlsx, Microsoft Excel; Supplemental Table 1). The experiments are named as the date_prefix_experiment number_protocol number. All cells recorded from the same animal share the same experimental date.

The current clamp data (see “Current Clamp” folder) contains two subfolders, “Step Protocol” and “Frozen Noise”. Step Protocol data includes two channels (voltage and current), each of which includes two columns (timestamp and voltage/current values in volt and amp, respectively) for each repetition. Users can visualize both the current injected to clamp the soma and the observed voltage response. Data from each stimulus condition is saved under a separate variable which starts with “Trace_a_b_c_d” and includes information about a) the cell and experiment ID, b) the data type, c) the number of sweeps in each dataset, and d) the channels.

The “Frozen Noise” subfolder contains the voltage trace (i.e. neuronal response to the injected frozen noise), hidden state (activity in the modeled network responds to, see [34] for details) and the injected current trace. In addition, a Matlab “struct” variable named “settings” is provided. Settings provide metadata under following “fields”: condition, experimenter, baseline (membrane potential value (in mV) at which the cell is kept with the baseline current injection), amplidute_scalind (the scaling factor used to translate the output of the neural network, in pA value), tau (the time constant that defines the average switching speed of the hidden state), mean_firing_rate (of the artificial neurons), sampling_rate (the acquisition rate (in kHz)), duration (in ms), FLAG_convert_to_amphere (a binary value that is 1 if the output was converted into Ampere), and cell_type (regular spiking vs fast spiking).

The voltage clamp folder includes two subfolders: VC Step (voltage step-and-hold) and VC Sawtooth, the latter containing 3 subfolders with recordings from experiments with triangular sweeps at 3 frequencies (5, 10 or 50 Hz). Data in the Voltage Clamp folder is organized similarly to data in the Current Clamp folder, and variable naming follows the formatting rules described above.

### Cell type classification

K-means clustering (cluster count=2; the number of repetition=10) was performed to classify neurons into fast-spiking and regular spiking neurons, using current clamp step-and-hold recordings. The clustering was based on the maximum firing rate reached during the current step injections and on the mean spike half-width across all stimulus steps during the current-clamp, step-and-hold protocol. Please note that the cell classification is solely provided to help the user to navigate the data. We do not claim that neurons can be necessarily electrically classified in a binary fashion, nor do we claim that commonly utilized clustering approaches are optimal for accurate (albeit broad) classification of excitatory (mostly regular spiking) and inhibitory (predominantly fast-spiking) neurons.

### Re-use potential

The dataset is rich in information regarding current versus voltage dynamics in adult cortical neurons. The independent variables in the database are the sex and age of the animal. While current-clamp experiments provide information about sub- and suprathreshold voltage dynamics, the voltage-clamp experiments are informative about the ionic conductances that lead to activation or inactivation of neurons.

In the step-and-hold current-clamp experiments, the voltage responses can be quantified using subthreshold (e.g. amplitude, latency, duration of the postsynaptic potential) and suprathreshold (e.g. interspike interval adaptation, spike count, spike half-width) responses to somatic current injection (Figure 3). Because multiple stimuli with incrementally increasing current intensities are delivered, cellular responses can be mapped onto stimulation intensities, allowing users to study input/output curves for the parameters of interest.

**Figure 3.**
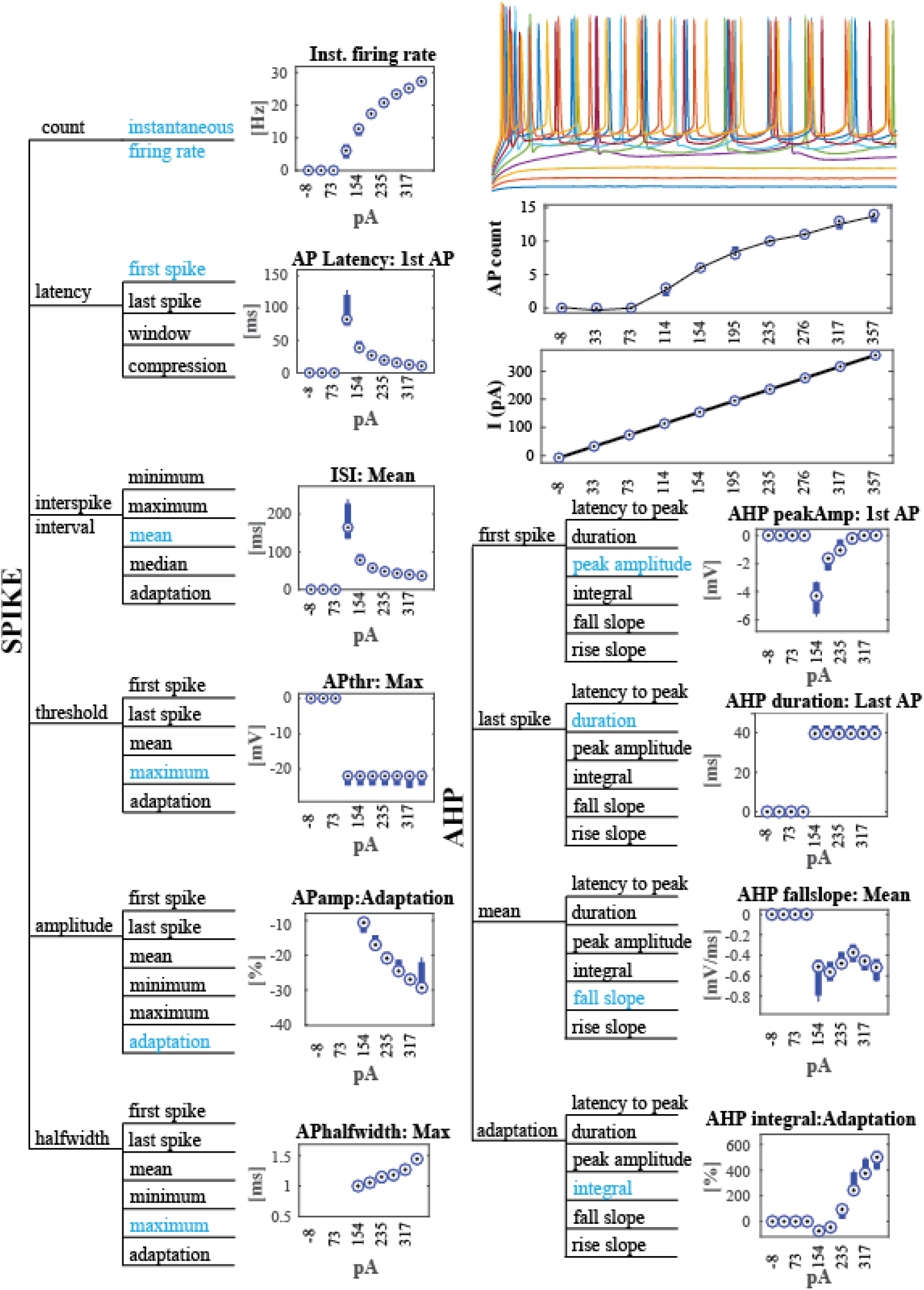
Electrical characterization of the spiking response in current-clamp experiments. The parameter space is shown as a hierarchical tree. Variables shown in blue are used for the data displays. AHP = afterhyperpolarization. “Compression” is a normalized metric that can be calculated as the difference between observations (e.g. spike timing) over the duration of stimulus. In the case of “latency compression” it is calculated as the temporal difference between the first and last action potential divided by the stimulus duration. “Adaptation” is the relative change in the observed variable, normalized to the first event. For example, in the case of spike amplitude adaptation, it is calculated as (AP_amp_first-AP_amp_last) / AP_amp_first. All characteristics are measured relative to the stimulus amplitude (the current injected, in pA). Membrane potential traces on the top-right are the responses to incremental current injections, superimposed on top of each other. The data below the raw traces represents the number of action potentials and the amplitude of the injected current across the 10 step-and-hold stimuli in this experiment (Filename: 170130_AL_133).

Action potentials can be studied both in terms of their shape (e.g. waveform, rise and decay slope, amplitude of the positive and negative peaks, the half-width of spike) and temporal response properties (that allow quantification of the rate and timing of action potentials during synaptic activation). Since adaptation to a sustained current injection is commonly used as a criterion to classify neurons, the data provides an inclusive database for the electrical classification of adult neurons, creating synergy with other publicly available databases, e.g. Neurodata Without Borders [36] and the Allen Institute Cell Type database [37]. The data can be used independently or in the context of computational models of neural networks, a broad selection of which can be found in the ModelDB database [38].

In addition to sustained somatic depolarization, the current-clamp database also includes “frozen noise” injections, during which a time-varying current was injected into the recorded neuron (Figure 4). The injected current was generated using an artificial neural network (see [34] for details) of 1000 neurons, each one firing spike trains from an inhomogeneous Poisson process, responding to a binary hidden state which represents the presence or absence of an external stimulus. The activity of all the neurons in the artificial network is integrated and the resulting current is corrected for the baseline current required to keep the patched neuron at −70mV. This summed current is injected to the patched soma. A major utility of the frozen noise protocol is that it allows direct quantification of neuronal information transfer [10, 34]. Compared to other metrics of neuronal information transfer [18, 39, 40], this approach enables bias-free quantification of information with a short (3 or 6 min) stimulation protocol [34].

**Figure 4.**
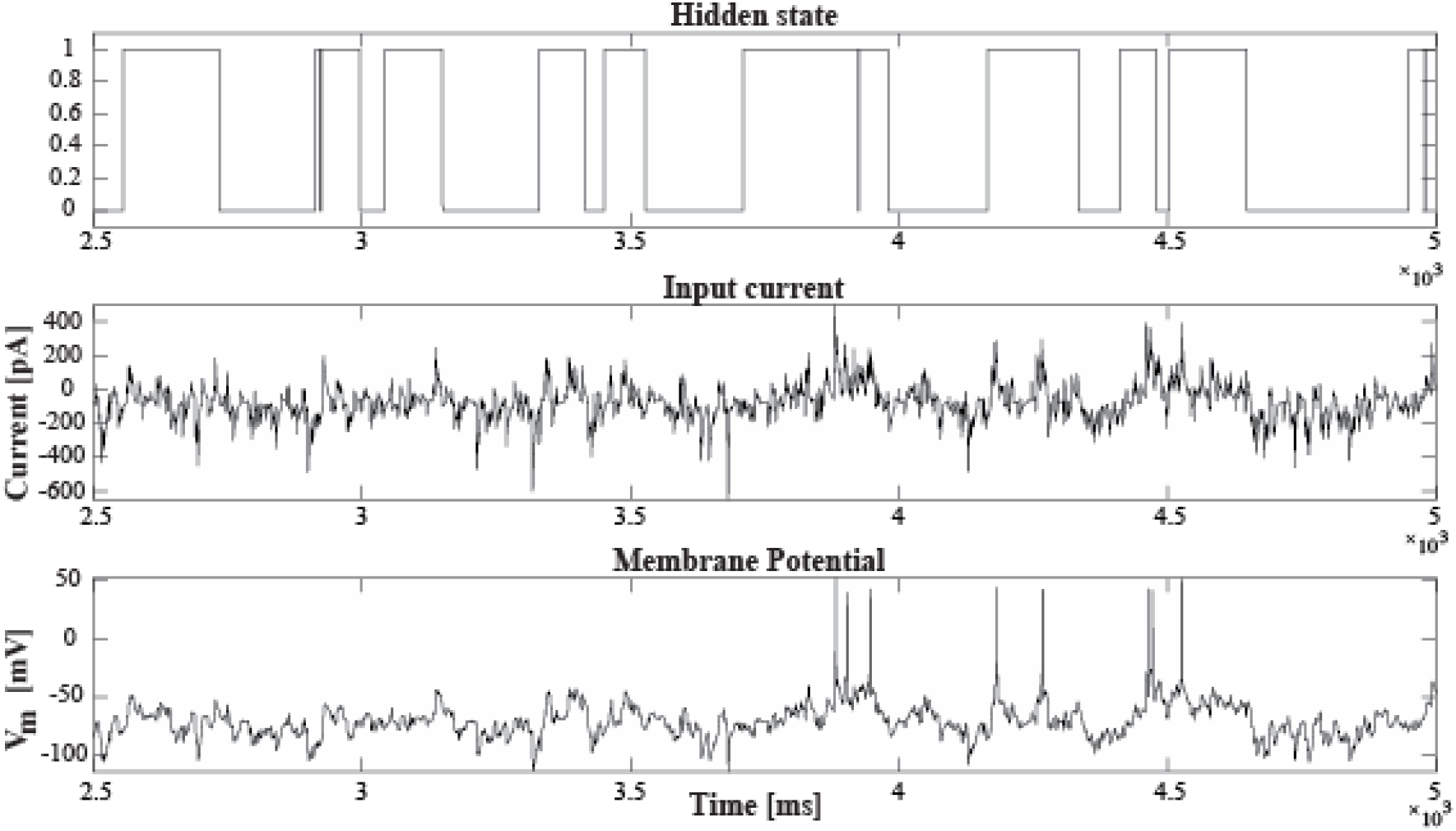
Frozen noise injection in current-clamp configuration. Representative recording from a single neuron (experiment 171207_NC_146). Top row: Binary representation of the hidden state that forms the input to an artificial neural network with 1000 point neurons, firing action potentials following an inhomogeneous Poisson process (see [34] for details). Middle row: the synaptic current generated by the artificial network that was injected into the recorded neuron. Bottom row: the membrane potential response of the recorded neuron.

In the database, experimental data recorded from our frozen noise protocol include the recorded membrane potential voltage, the hidden state and the current injected into the neurons (Figure 4). Thus, the user can perform forward and reverse modeling to predict the neuronal response and to study neuronal dynamics in the adult neocortex.

Going beyond the voltage dynamics in the adult neurons, the database also provides insight into the ionic currents that flow through the membrane. With the triangle shaped VC-Saw protocol (Figure 5), it is possible to measure the activation threshold of the currents flowing through the membrane, by assessing when deviations from the expected sawtooth shape occur. Additionally, it is possible to compute: amplitudes and latencies of the events, peaks half-widths, the percentage difference between consecutive events and the total number of events in each sweep. Other features could also be extracted from the dataset depending on the researchers’ interests.

**Figure 5.**
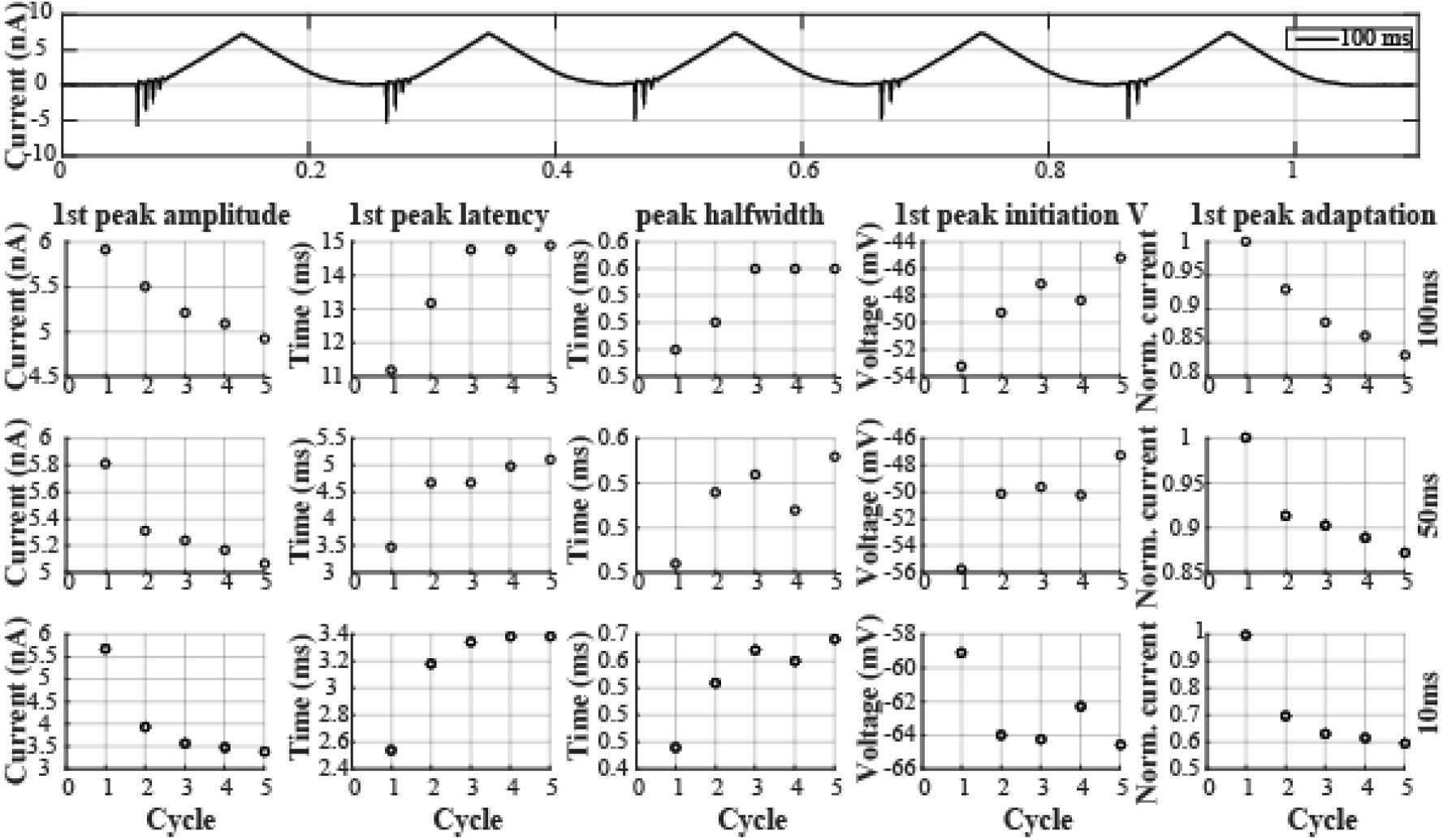
Voltage-clamp sawtooth protocol. Top row: Current trace from a representative experiment (180412_AB_53_ST). Figurines, left to right, are measurements of the first peak amplitude, first peak latency, the half width of the first inward current, membrane potential at which the inward current is initiated, and the adaptation of the first event amplitude across the 5 (triangle) cycles. Data in the bottom three rows are from three different sawtooth speeds (10/50/100 ms, corresponding to 100/20/10 Hz stimulation). The five points in each figurine are calculated from the first inward current in each (triangle) cycle.

The current-voltage relationship was measured with voltage-clamp steps (Figure 6), which could be used to produce an I/V curve. The peak amplitude, latency, and peak half width can be extracted for the inward currents observed during the sustained depolarization of the soma.

**Figure 6.**
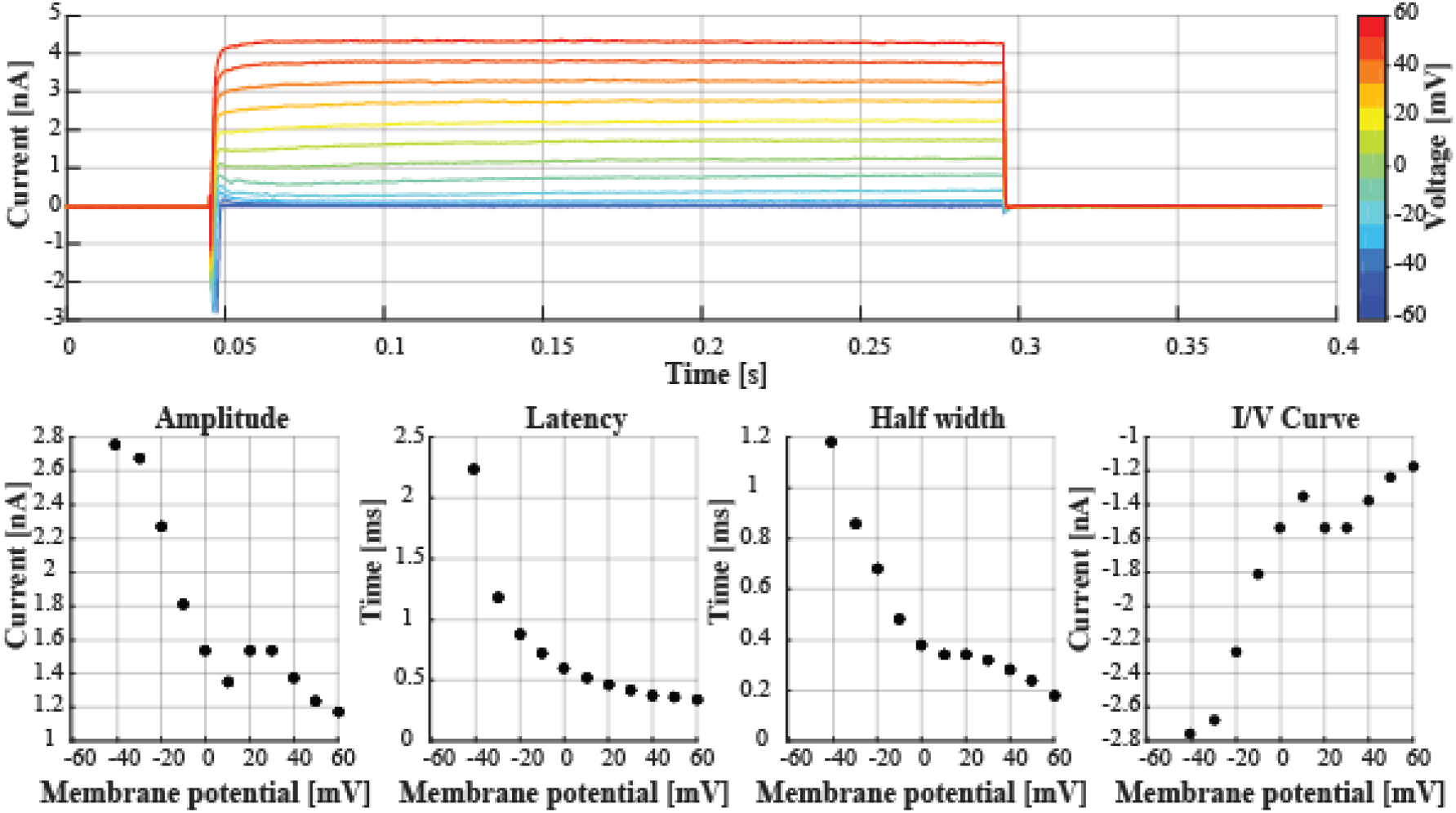
Step-and-hold protocol in voltage-clamp preparation. Top panel shows data from a representative experiment (170915_AB_5_VC). Every other figurine shows one of the analyzed features, including the amplitude of the peak, temporal delay between the stimulus onset and the peak amplitude (i.e. latency), width of the evoked transient measured at half maximum as well as the current at every holding potential (I/V curve).

### Application scenarios

From the recordings available in this database, it is possible to quantify active the membrane properties of supragranular layer neurons to infer current and voltage dynamics during somatic depolarization.

Network development is based on processes of self-organization that are highly dependent on sensory stimuli and experience [2]. Such plasticity is not limited to early life development. From the voltage-clamp and current-clamp experiments, it is possible to infer the biophysical properties of layer 2/3 pyramidal neurons of the adult somatosensory cortex under baseline conditions, in the absence of altered sensory experience.

The focus on adult neurons brings a new perspective to the study of membrane properties, as data from this mature ages are still scarce. The dynamics of the active electrical properties of the membrane can be accessed as a function of different developmental time points and/or sex, and the recorded data can be used as virtual neurons in dynamic-clamp experiments.

In a computational approach, spiking properties described herein could be used for biomimetic modeling of diverse networks, facilitating the study of computational roles of circuit motives. Moreover, applying the principles of information transfer and recovery to the data might help recreate neuronal functions in artificial systems.

### Limitations

Neurons in this dataset originate from regular spiking and fast spiking neurons, however, there is no anatomical characterization of the neuron type studied. The database is focused on Layer 2/3 of the somatosensory cortex as a model region and does not allow the study of neuronal information processing across different cortical regions in isolation. However, the user might consider comparing data across different regions and species by utilizing the other publicly available databases, e.g. Neurodata Without Borders [36], the Cell Type database [37] of the Allen Institute and the Collaborative Research in Computational Neuroscience data sharing initiative [41].

## Acknowledgements

We would like to thank the members of the Department of Neurophysiology, Donders Institute for Brain, Cognition and Behaviour for stimulating discussions and the critical insight on the manuscript.

